# Methods to assess neuronal primary cilia electrochemical signaling

**DOI:** 10.1101/2025.04.01.646689

**Authors:** Paul G. DeCaen, Louise F. Kimura

## Abstract

Primary cilia are polymodal sensory organelles which project from the apical side of polarized cells. They are found in all brain hemispheres but are most pronounced in neurons which comprise the granular layers of the hippocampus and cerebellum. Pathogenic variants in genes which encode primary cilia components are responsible for neuronal ciliopathies— a group of central nervous system disorders characterized by neurodevelopmental conditions such as intellectual disability, seizure, ataxia, and sensory deficits. In the hippocampus, neuronal primary cilia form chemical synapses with axons and their membranes are populated with unique sets of ion channels and G protein-coupled receptors (GPCRs). Primary cilia are small and privileged compartments that are challenging organelles to study. In detail, we describe cilia electrophysiology methods and the use of cilia-specific fluorescent sensors to assay neuronal polycystin channel function and serotonergic receptor signaling, respectively. These tools allow researchers to assay calcium, cAMP and channel-related signaling pathways in isolated neurons in real time and in semi-quantitative terms, while enhancing our understanding of this understudied organelle and its dysregulation in ciliopathy disease states.

## 1 Introduction

Neuronal primary cilia act as specialized sensory and signaling hubs— receiving and processing signals from the external environment to coordinate unique forms of electrochemical communication^1-3^. Ciliogenesis in most types of neurons occurs during G0 and G1 phases of the cell cycle, utilizing mitotic microtubule machinery which are dynamically regulated^4^. The mother centriole migrates to the apical surface of the neuron, forming the basal body of the primary cilium and the nucleation site of microtubules of the internal scaffolding structure known as the axoneme array (9+0). Once fully developed, the primary cilium extends 2–12 μm from the apical surface of the neuronal plasma membrane, with a diameter of less than 500 nm^5-8^. At the neuronal primary cilia-cell junction, the basal body partners with the BBSome— a protein complex which, along with diffusion barriers present at the ciliary base, give the organelle its privileges status and unique composition. Once inside the cilia, the intraflagellar transport (IFT) system shuttles ciliary proteins and other cargo along the axoneme via motor proteins^9,10^. Here, critical cilia signaling pathways such as Hedgehog and Wnt that are vital for maintaining neuronal polarity, cell fate determination, and tissue patterning are established^11-14^. Primary cilia from hippocampal neurons form serotonergic synapses with axons on brain stem neurons (axo-ciliary synapse), which modulates the post synaptic neuron’s epigenetic state through Gαq11-Rho mediated chromatin accessibility changes^15^. In addition, neuronal primary cilia in disparate locations within the central nervous system express unique GPCRs, such as SSTR3 (cerebral cortex, hippocampus, hypothalamus, cerebellum, amygdala), MC4R (hypothalamus, brainstem) and 5-HT6R (hippocampus, cerebral cortex, striatum)^16-18^. In various regions within the CNS, SSTR3 is coupled to Gαi, while MC4R signals through Gαs and crosstalks with other ciliary proteins such as ADCY3^19,20^. Whereas 5-HT6R signals through Gαs but also promotes Gαq11-Rho mediated signaling^15,20-22^. These cilia-specific signals regulate a wide range of physiological functions from appetite control; metabolic regulation; cognitive processes and sensory perception. Disruption of GPCRs and channels in the cilium can lead to significant brain developmental disorders and animal neurophenotypes, underscoring their critical role in neuronal function^23,24^.

The importance of the primary cilia in human health is highlighted by more than 35 congenital diseases called “ciliopathies”, which impact the development of organ systems such as the kidney, brain, heart and eye^25,26^. A subset of these syndromes can be classified as neuronal ciliopathies as the gene variants cause defects in the structure or function of primary cilia in neurons (**Table 1**)^23,27^. For example, Joubert syndrome and Bardet-Biedl syndrome are caused by variants in the genes encoding for cilia assembly components— such as centrosomal CEP120 and BBS proteins, and the GTPase ARL13B^28^. These syndromic diseases primarily impact the development of the central nervous system but also have comorbidities affecting organ systems outside of the brain. Developmental defects are common amongst individuals with neuronal ciliopathies, such as polydactyly (extra digits on hands and toes) and *situs inversus* or *totalis* (inverted positioning of body organs)^29^. In addition, common comorbidities shared by neuronal ciliopathies are polycystic kidney disease (PKD) and other renal defects such as nephronophthisis, but the reason for this is not known^26,30^. However, since the autosomal dominant form of PKD (ADPKD) is caused by gene variants in renal cilia calcium channel subunits (PKD1 and PKD2), and since other ciliopathy variants impact Ca^2+^ effectors (e.g. CEP 290) share renal cyst comorbidities— localized ciliary Ca^2+^ dysregulation might be a unifying disease-causing mechanism observed in these conditions^31-35^.

**Table 1:** Neuronal ciliopathy syndromes. A tabulated list of neuronal ciliopathies with their associated genetic variants and clinical pathology. Note some neuronal ciliopathies have different diagnoses but are associated with variants that impact the same gene. Renal abnormalities comorbid with neuronal ciliopathies most frequently manifest as early onset polycystic kidney disease.

| Syndrome<br>(Gene variants associated) | Pathology |
| --- | --- |
| **Joubert syndrome** (*CEP290; TMEM67; AHI1; CC2D2A; INPP5E; ARL13B; RPGRIP1L; KIF7; TCTN1-3; MKS1; TMEM216*) | Characterized by abnormal brain development, leading to motor and cognitive impairment. |
| **Bardet-Biedl syndrome** (*BBS1-18 includes MSKKS, LZTFL1, SDCCAG8 and ARL6; IFT27*) | This multisystem disorder often involves cognitive impairment, obesity, and vision loss due to retinal degeneration. |
| **Meckel-Gruber syndrome** (*MKS1; TMEM67; CC2D2A; TMEM216; CEP290; B9D1; TMEM231; RPGRIP1L ; TCTN1-3; TMEM107; NPHP3*) | A severe disorder that often leads to brain malformations and is usually lethal early in life. |
| **Alström syndrome** (*KIF7; WDR35; DYNC2H1; IFT80; MKS1*) | A condition that can cause vision and hearing loss, obesity, and heart failure, along with neurological symptoms. |

Our understanding of the primary cilia in neurons has been restricted by the limitations of traditional tools to assess dynamic chemical changes within this privileged organelle. Methodologies such as immunofluorescence of fixed samples only capture static changes in cilia signaling. Neuronal primary cilia do not load with standard calcium dyes (e.g., Fura) or cAMP fluorescent reporters (e.g., unconjugated cADDis) where their diffusion is limited to the soma, axon, and dendritic compartments. To address this, several groups have developed cilia-localized ratiometric Ca^2+^ sensors, and reporters that enhance fluorescence intensity in response to ciliary cAMP concentration (**Table 2**)^36-38^. Here, we describe methods for the deployment of Ca^2+^ and cAMP sensors, both in isolation and in combination, to assess cell signaling in primary cilia of cultured hippocampal neurons. Although it is established that neuronal primary cilia are richly populated with polycystin transient receptor potential (TRP) channels encoded by PKD2L1— their ion channel activity cannot be assessed by voltage clamping the soma or cell body membranes^39^. To address this, we provide experiment details to directly record neuronal primary cilia PKD2L1 channels aided by the expression of genetically encoded fluorescent reporters (**Table 2**)^6,40-42^. Outcomes proved a methodological toolbox to studying ciliary chemical and electrical activity in real time to understand the basic biology of this understudied organelle and its dysregulation in ciliopathy disease states.

**Table 2.**
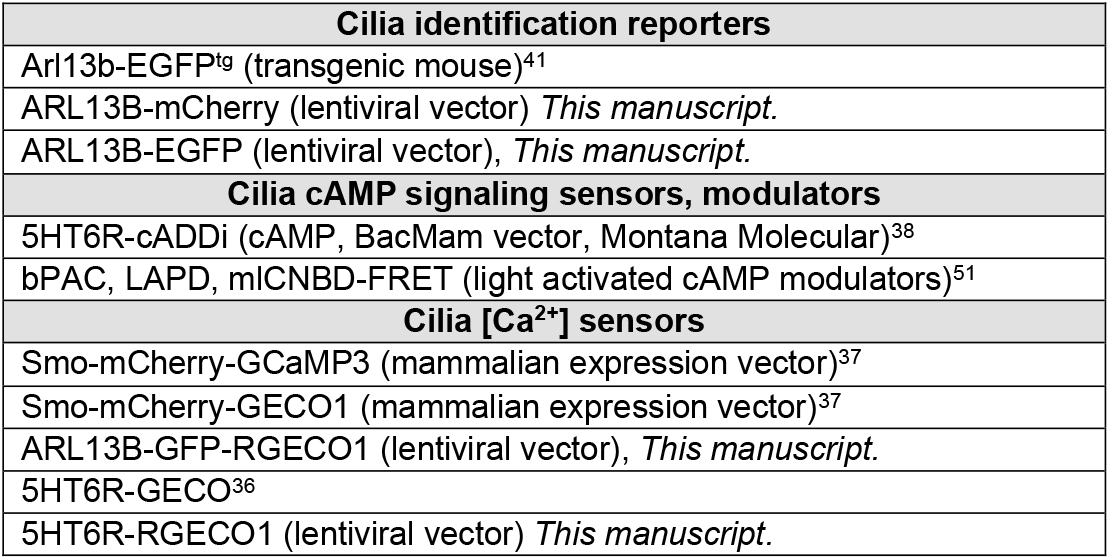
A primary cilia toolbox. Genetically encoded fluorescent reporters for identification, Ca^2+^ concentration and GPCR signaling within primary cilia. Previously published expression constructs or transgenic animal models are listed with corresponding citations.

## 2 Material and methods

### 2.1 Hippocampal neurons primary culture

All mice utilized were housed in AAALAC-approved Center for Comparative Medicine (CCM) at Northwestern University, Feinberg School of Medicine. All procedures with mice were in accordance with the recommendations of the Panel on Euthanasia of the American Veterinary Medical Association. Animals were maintained in temperature-controlled rooms on a 12 h day/light cycle, fed *ad libitum*, and cages are changed 2-3 times weekly. Wt or stable expression of the ARL13B transgene conjugated to enhanced green fluorescent protein (*Arl13B-EGFP*^*tg*^) in the genome of a founder C57BL/6 mouse were used. Importantly, *Arl13B-EGFP*^*tg*^ mice do not have a distinct behavioral or anatomical phenotype from C57BL/6 mice^38,45^. Hippocampi were dissected from P0-P1 mice (n= 6-8 pups per experiment) under a scope in a cold solution Hank’s Balanced Salt Solution (HBSS, 14185-052, Gibco) containing HEPES (100 mM) (15630-080, Gibco) and digested in Trypsin 0.025% (25-050-CI, Corning) at 37°C for 5 minutes. Solution was removed and the tissue were gently triturated using a P1000 in 1 mL of platting media which consisted of Neurobasal A (10888-022, Gibco) containing B27 supplement (17504044, Gibco), 1 % penicillin/streptomycin (15140148, Gibco), 10 % normal horse serum (NHS, 260500, Gibco), 1% glucose (A2494001, Gibco) and GlutaMAX (350500-061, Gibco). Cells were seeded only in the glassed surface area of the dish/coverslip using platting media at 1 x 10^6^ cells/dish or coverslip. Glass bottom dishes (P35G-1.0-14-C, Mattek) or coverslips were coated only on the glassed surface area using Matrigel (1:20, 356234, Corning). Media was replaced with a feeding medium (media without NHS) after 3 hours of seeding. After 1-3 days in culture, WT hippocampal neuron cultures were infected with lentivirus vector encoding for cilia reporters and sensors (See section 2.3). All hippocampal neurons (WT or *Arl13B-EGFP*^*tg*^) were cultured for up to 14 days and the feeding medium was changed every 2-3 days.

### 2.2 Electrophysiology

Below are described strategies for measuring ciliary ion channels from cells expressing cilia-specific fluorescent proteins with “on-cilia” patch clamp configuration (**Figure 1A**). This method is an extension of the voltage clamp method established more than 50 years ago with details optimized for cilia membrane ion channel recordings^6,41,43^. Electrophysiology experiments were performed using either an inverted confocal or widefield microscope (**Table 3**). We used a 20x objective lens to locate cells projecting the primary cilium above the focal plane or to the side of the cell, and having sufficient fluorescence, then switched to a 60x water emersion objective with a 2x photomultiplier to visualize the cilia during seal formation. To capture images and video recordings of the cilia patch configuration, a standard CCD or CMOS camera on a widefield microscope sufficient.

**Table 3.** List of equipment used with manufacture part numbers.

| COMPONENT | PART NUMBER, MODEL | MANUFACTURE |
| --- | --- | --- |
| <u>Bright field microscope components</u> |  |  |
| Equipped with standard components | IX73 | Olympus |
| Objective 20x, Air U Plan Fluorite, NA 0.5, WD 1.6 mm | 1-U2B525 | Olympus |
| Objective 40x, Air U Plan Fluorite, NA 0.75, WD 0.51 mm | 1-U2B527 | Olympus |
| Objective 60x, water emersion NA 1.2, WD 0.51 mm | UPLSAPO | Olympus |
| Fluorescent light source | Lumen 200 | Prior |
| Filter for EGFP, GCaMP3, mEmerald 470/20x, 485DM, 517/45m | U-FGFP BX3 GFP/BLUE WIDE | Olympus |
| Filter for EYFP, Venus, Citrine 495/10x, 505DM, 537/45m | U-FYFP; BX3 YFP/YELLOW NARROW | Olympus |
| Filter for RFP, mCherry, DsRED 545/20x, 565DM, 597/55m | U-FRFP; BX3 RFP/GREEN WIDE | Olympus |
| **Electrophysiology and imaging components** |  |  |
| Patch clamp amplifier | Axopatch 200B | Molecular Devices |
| Digital to analogue signal converter | Digidata 1440 | Molecular Devices |
| CMOS camera | ORCA-FLASH 4.0LT | Hamamatsu |
| CCD camera | U-TV1X-2-7 | Olympus |
| Rapid perfusion system | VC-77CS "Fast step system" | Warner Instruments |
| High speed pressure clamp system | HSPC-1 | ALA Scientific Instruments |
| Dual micromanipulator System (Stepper motor driven) | MP-300 | Sutter Instruments |
| Dual micromanipulator System (Linear piezo driven) | Inchworm, uM 3-axis System 69-3432 | Sensapex |
| **Microelectrode fabrication components** |  |  |
| Flaming brown filament type micropipette electrode puller | P-97 3 mm box filament Settings: Heat = ramp test value; Pull = 44; Velocity = 65; Delay= 200; Pressure = 600 | Sutter Instruments |
| Laser-Based micropipette electrode puller | P-2000 | Sutter Instruments |
| Capillary glass O.D. 1.5 mm, I.D. 0.84 mm | 1B150F-4 | World Precision Instruments |

**Figure 1:**
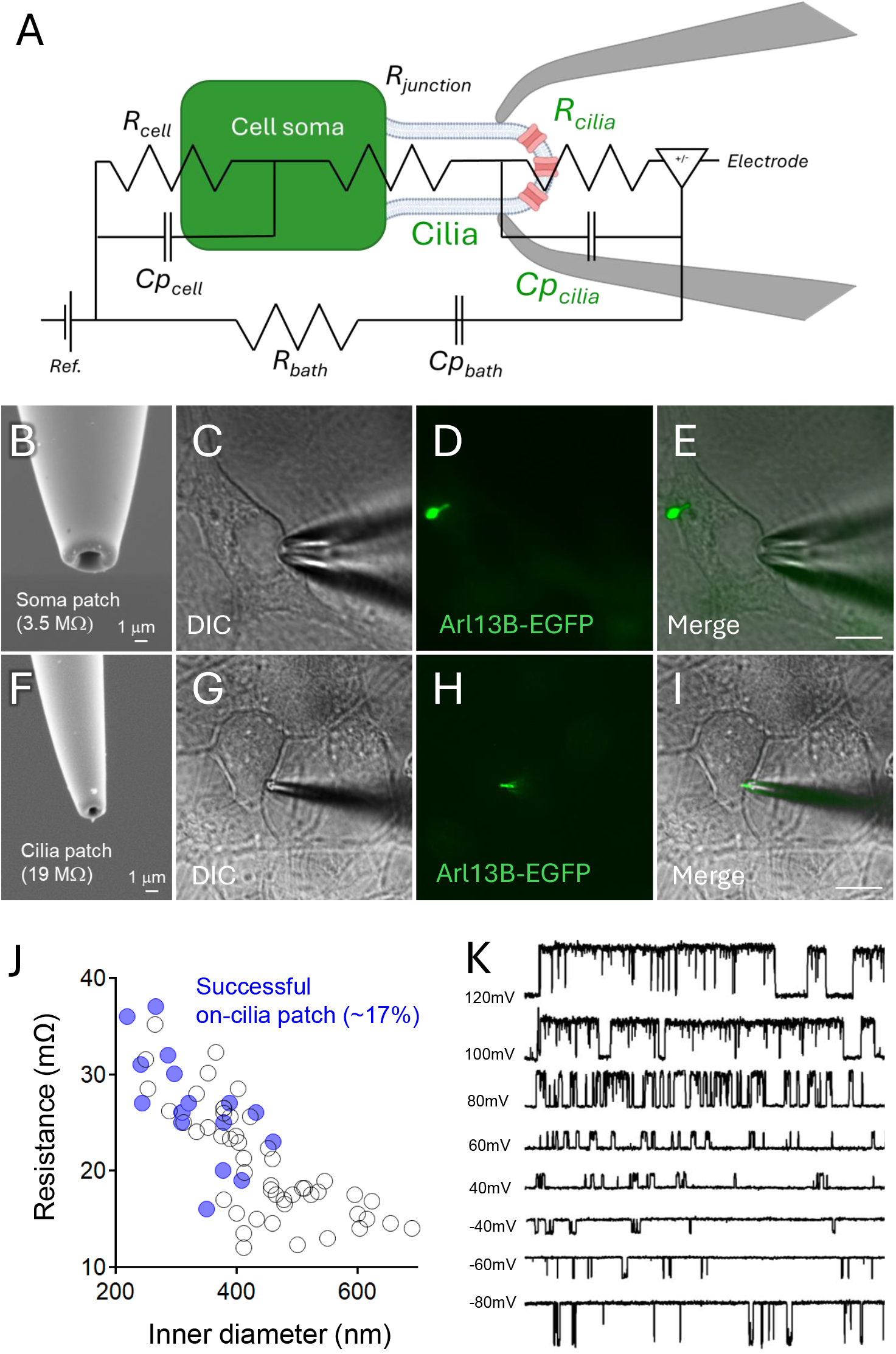
Neuronal primary cilia electrophysiology. We used the “on-cilia” patch clamp configuration with equivalency circuit as represented (**A**). Hippocampi was dissected from P0 mice expressing cilia-specific fluorescent protein (Arl13B-EGFP^tg^ mice) and cells were seeded in glass bottom dishes. Ciliated neurons were patched (DIV 6-10) using the whole cell (**B-E**) or cilia patch clamp (**F-I**) configurations. Scanning electron microscope images of a whole-cell (**B**) and cilia patch (**F**) electrodes were taken for comparison purposes between the two techniques. The correlation between electrode resistance versus its lumen diameter with successful giga-seal formation (blue circles indicate successful trials) is represented in Figure 1J. The current traces of PKD2L1 channels present in the cilium are represented in Figure 1K. Scale bar for E, I: 10 μm.

#### 2.2.1 Glass electrode fabrication

Filamented glass with inner and outer diameters of 0.84 and 1.4 mm, respectively were used. The resistance of the “pulled” electrodes manufactured using a laser-heated puller were between 5-9 MΩ with an inner aperture of 0.5-1 μm in diameter. These electrodes were then polished using a commercially available electrode polisher (**Table 3**). This step smoothens the surface of the glass and narrows the aperture of the tip of the electrode, thereby increasing the resistance of electrode. To determine the optimal pipette resistance for the cilia recording, the resistance capable of achieving a high resistance seal within the cilia membrane (>10 GΩ) were correlated from many attempts. Afterwards, we used a scanning electron microscope to measure the inner diameter of each patch electrode used.

#### 2.2.2 Single channel recordings

Single channel currents were measured from the primary cilia membrane an Axon 200B (Molecular Devices) amplifier connected to Digitdata 1550 digitizer (**Table 3**), as previously described^6,41^. The standard pipette solution contained (in mM): 100 NaCl, 10 HEPES, pH 7.4 with NaOH and adjusted to 300-305 mOsm/L with mannitol. High potassium bath solutions containing 130 KCl, 15 NaCl, 10 HEPES, 1.8 CaCl_2_, 1 MgCl_2_, pH 7.4 with KOH and 300-305 mOsm/L, were used to neutralize the membrane potential. Of note, the composition of the saline which determines the ion to be used as a charge carrier depends on the type of ciliary ion channel opening events to be tested. To measure the PKD2L1 channels activity on the neuronal cilia membrane, as exampled here, the pipette solution contained (in mM): 100 CaCl_2_, 10 HEPES, pH 7.4 with Ca(OH)_2_ and adjusted to 300-305 mOsm/L with mannitol. The permeability of monovalent cations (P_X+_/P_Na_) can be determined by observing the shift in reversal potential when the external Na^+^ bath solution is replaced. Recording saline was front filled and backfilled into these pipettes to ensure electrical continuity. Data was collected using an Axopatch 200B patch clamp amplifier and the pClamp 10 software (Molecular Devices, **Table 3**). As a rule of thumb, we recommend sampling the current 5x more frequently (25 KHz) than the low-pass Bessel filter rate. While this makes the size of the acquisition file larger, these settings allow for the sufficient sampling of both single and whole cilia currents, which can be re-sampled during the analysis phase.

### 2.3 Cilia-targeted identification reporters, Ca^2+^ sensors and cAMP sensors

Cilia-targeted identification reporters (ARL13B-EGFP; ARL13B-mCherry); Ca^2+^ sensors (ARL13B-GFP-RGECO1; 5-HT6R-RGECO1) were custom synthesized (VectorBuilder) into the lentiviral cassettes (pLV[Exp]-puro-CMV) to generate lentivirus (10^8 titer) solutions encoding for each reporter or sensor. Primary hippocampal neurons isolated from WT or ARL13B-EGFP^tg^ mice were transduced overnight at DIV 2-3 (MOI=1-2) in a solution containing feeding media and polybrene (4 μg/mL). Experiments we carried out from 3 days after transduction. For the cAMP sensor coupled to the 5-HT6R encoded by the BacMam vector (cilia-targeted cADDis sensor, D0201G, Montana Molecular), cultured neurons from WT mice were used and transduced overnight at DIV 5 according to the manufacturer instruction. Fluorescent intensity of the cADDis (Green) decreases in response to increases in cAMP. Experiments we conducted from 1 day after transduction. When both 5-HT6R-RGECO1 and 5HT6R-cADDis sensors were evaluated together, cells were transduced as mentioned above with cADDis transduction performed one day prior to the experiments. Ciliated neurons were filmed (1 frame/sec) before and after the application of different solutions: To calibrate the ARL13B-GFP-RGECO1 sensor, different calcium concentrations (0, 1, 10 or 100 μM) in the presence of ionomycin (1 μM) were applied. Bath standard solution was composed by (in mM): 140 NaCl, 10 HEPES, 5 KCl, or otherwise stated, and pH 7.4 was adjusted with NaOH. Ratiometic fluorescence intensity was calculated by the ratio of RGEGO1/GFP and normalized by the fluorescence average of the initial 30 sec of filming. For the 5-HT6R-RGECO1 sensor alone, solutions containing ionomycin alone (1 μM) or with calcium (10 μM) were assessed. For the cADDis alone, the 5-HT6R agonist WAY 181187 (1 μM) or in a combination with the 5-HT6R antagonist SB 271046 (100 μM, pre-incubation for 10 minutes) was tested. When neurons were transduced with both 5-HT6R-RGECO1 and cADDis, WAY 181187 followed by ionomycin (1 μM) + Ca^2+^ (10 μM) were applied. Fluorescence intensity was calculated by the ratio of RGEGO1 (red)/GFP (green) and normalized by the fluorescence average of the initial 30 sec of filming, or by the ΔF/F0 in which F0 is the fluorescence average of the initial 30 sec of filming.

### 2.4 Drugs and salts

Salts used to prepare bath solutions are manufactured by Sigma Aldrich (St. Louis, MO) or Fisher (Waltham, MA). Ionomycin free acid (ref # 407950, Millipore Sigma), WAY 181187 oxalate (ref # 5589, Tocris Biosciences, UK), SB 271046 hydrochloride (ref #3368, Tocris Biosciences, UK) and thapsigargin (ref # T7458, ThermoFisher) were initially dissolved in DMSO (ref # D2650, Sigma Aldrich) and diluted in working solutions according to each experiment. α-hemolysin (ref #H9395, Sigma Aldrich) was dissolved in water (stock solution of 0.5 mg/mL).

### 2.5 Data analysis

Electrophysiology data was analyzed with ClampFit (Molecular Devices) and IGOR Pro (Wavemetrics) software. Data collected with the sensors were evaluated for normality using Anderson-Darling test. For Arl13B-GFP-RGECO1 ratiometric experiments, one-way ANOVA followed by Holm-Sidak’s multiple comparisons test were used to analyze the data of peak intensity. For the 5-HT6R-RGECO1 experiments, multiple paired t-tests were used.

## 3 Results

### 3.1 Neuronal primary cilia electrophysiology

Glass electrodes with submicron aperture are fabricated to establish high resistance seals with the primary cilia membrane to measure stochastic single channel open events (“on-cilia” configuration) or total ciliary current (“whole-cilia” configuration)^40,41^. After forming giga-ohm resistance seals, the neuronal cilia can be severed (“inside-out” configuration) from the soma to access and apply effectors to the internal side of the cilia membrane (**Figure 1A-F**)^41^. The primary cilium is electrically insulated from the cell body, with its own resting membrane potential (−17 mV) and Ca^2+^ calcium concentration (350-700 nM)^36,37,41^. To match the geometry of the neuronal primary cilia, microelectrodes are fabricated in two steps. First, long taper borosilicate electrodes with 900-600 nm tip diameters are created using a heated filament or laser glass puller. Second, the electrode tip is polished using a microforge so that the lumen approximates the diameter (∽350 nm) of the primary cilia (**Figure 1B, E**). Neuronal primary cilia can be visualized using brightfield microscopy, but to faithfully distinguish these structures from filipodia, we used genetically encoded fluorescent ciliary markers (**Table 2**). Here, the cilia specific proteins such ARL13B are conjugated with fluorescent protein and expressed as a transgene in the mouse genome (ARL13B-EGFP^tg^) or transduced virally in cultures of neonatal hippocampal neurons (ARL13B-GFP or ARL13B-mCherry). We observed that cultured for 6-10 days offered the highest number of ciliated neurons for our electrophysiology experiments (**Supplemental Figure 1**). We recommend selecting neurons with primary cilia projecting above the soma’s focal plane and towards the recording electrodes path of approach. The electrodes approach and course position of the ciliated neurons can be achieved with lower magnification (20-40x objective). However, once the recording electrode approaches 100 μm from the cilia, the objective magnification should be increased (60-100x water or oil) which have limited working distances. Turning off the fluorescent light source to minimize photobleaching of the cilium. Typical gear-driven manipulators can be used to patch primary cilia, but we recommend using piezo manipulators which have minimal drift (**Table 3**). Inward suction (2-10 mmHg) the tip of the cilia membrane seals onto the pipette electrode tip, creating electrical resistance (R_cilia_) greater than 10 GΩ. We have determined that electrodes with 280-410 nm inner diameter and resistance of 17-32 MΩ frequently achieved high resistance seals of ciliary membranes allowing the recording of channels activity (**Figure 1F-J**). Exemplar PKD2L1 single channel open events recorded from a hippocampal primary cilium in the “on-cilia” configuration are shown in **Figure 1K**. Importantly, the command potential and the current directionality are reported in the opposite polarity and inverted directionality in the “on-cilia” configuration, to be consistent with conventions for reporting electrophysiology data sets. Single channel open probability can be reliably determined over a series of 1-5 second epochs and respective unitary conductance can be estimated by fitting the average current magnitude over applied membrane potentials to fit a linear equation.

### 3.2 Calcium sensor Arl13B-GFP-RGECO1 in hippocampal neurons

Primary cilia have 5-fold higher resting Ca^2+^ concentration and contain calcium-binding proteins with high disassociation constants which are frequently impacted by genetic variants implicated in neuronal ciliopathies^36,37^. Because they are resistant to chemical loading by dye indicators, native neuronal cilia proteins (Smo, ARL13B, 5-HTR6) can be linked to calcium sensor fluorescent reporters (GCaMP, GECO). An additional non-calcium dependent fluorescent reporter (mCherry, EGFP) can be added to the sensor to normalize the Ca^2+^ fluorescence response and provide ratiometric measurements which allow quantification of free Ca^2+^ concentration within the primary cilia. For expression in neonatal hippocampal neuron cilia, we linked ARL13B-GFP with red-shifted RGECO1 (genetically encoded Ca^2+^ indicator for optical imaging) within a lentivirus expression vector. After 5-7 days in culture, murine hippocampal neurons become electrically excitable, generating spontaneous action potentials when assessed by current clamp^44^. At this time point, spontaneous primary cilia calcium dynamics are captured by the ARL13B-GFP-RGECO1 sensor transduced hippocampal neurons (**Figure 2A**). Thess endogenous calcium waves can be visualized in real time (1 frame/sec) using a three-dimensional structured illumination microscopy (3D-SIM) in standard solution containing 1.8 mM calcium, **Supplemental Video 1**). To minimize potential biases associated with external calcium transients, we conducted our experiments using a calcium-free standard bath solution, in which spontaneous calcium waves in the neuronal primary cilia were markedly diminished (**Figure 2B**). After exchanging external conditions in the presence of the Ca^2+^ ionophore ionomycin, we observed external Ca^2+^ dose-dependent (1-100 μM) increases in RGECO1 fluorescence intensity (**Figure 3A, Supplemental Video 2**). Ionomycin alone (no external Ca^2+^) increased cilia Ca^2+^-dependent fluorescence, suggesting that the ionophore is stimulating release of intracellular store Ca^2+^, which propagates into this cilia organelle compartment. Although not significant, when ionomycin was combined with 1 μM Ca^2+^, it was less effective at stimulating cilia RGECO responses compared to ionomycin alone. We speculate that ionomycin efficacy in triggering intracellular Ca^2+^ store release was impacted by its 1:1 binding ratio to free external calcium^45^. To verify whether intracellular calcium released from cytoplasmic stores was entering the primary cilia, we re-analyzed the data with ROIs on both base and tip of the cilia. In every neuron tested, ionomycin-induced calcium responses initiated from the cytoplasm and propagated to the primary cilium (**Supplemental Figures 2A, B**). In addition, we repeated the experiment but first applied thapsigargin (SERCA pump inhibitor) for depletion of ER Ca^2+^ stores of the neurons for 3 minutes. We observed that thapsigargin triggered a mild increase in cilia Ca^2+^, but an attenuated cilia Ca^2+^ response from externally applied ionomycin (**Supplemental Figure 2C**). These observations demonstrate the new primary cilia Ca^2+^ sensors sensitivity to external and store operated calcium signaling in hippocampal neurons. Moreover, the sensor also detected calcium transients under micromolar concentrations of external calcium (500 nM) after permeabilization with alpha hemolysin (10 μg/mL) for 3 minutes (**Supplemental Figure 2D**). Thus, the use of calcium-sensors such as ARL13B-GFP-RGECO1, can be deployed to further examine the relationship between store operated Ca^2+^ signals and neuronal primary cilia responses in the central nervous system.

**Figure 2:**
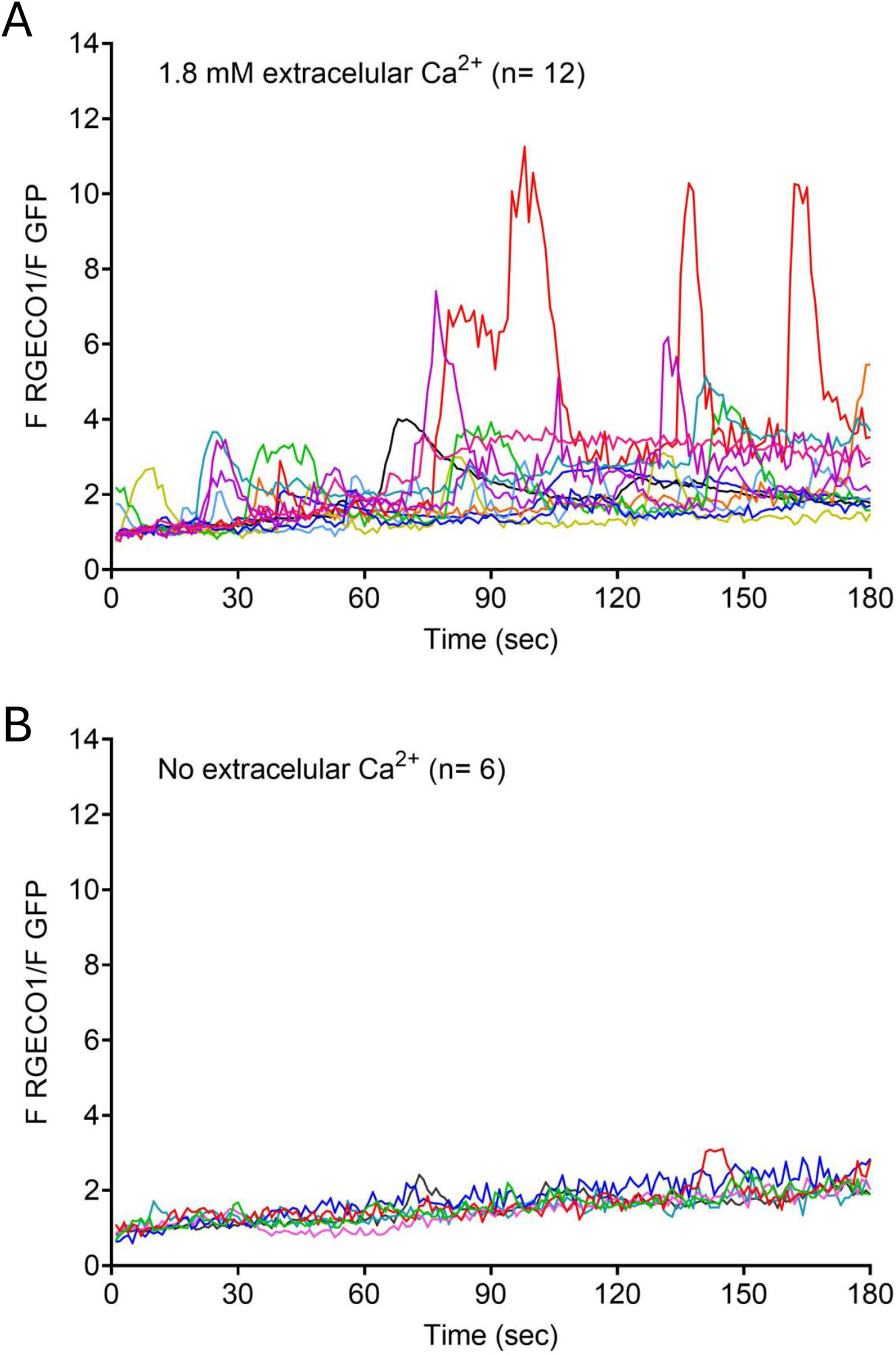
Spontaneous Ca^2+^ waves in the neuronal primary cilia. On DIV 2-3, hippocampal WT cultured cells were transduced overnight with the lentivector ARL13B-GFP-RGECO1 (MOI= 1-2) and 5 days after transduction, ratiometric experiments were performed under a confocal microscope. Neurons were imaged in the standard solution containing 1.8 mM of calcium (**A**) or in the standard solution (calcium-free) (**B**). Fluorescence intensity was calculated by the ratio of RGEGO1 (red)/GFP (green) and normalized by the fluorescence average of the initial 10 sec of filming, or when cells fluorescence returned to the baseline levels.

**Figure 3:**
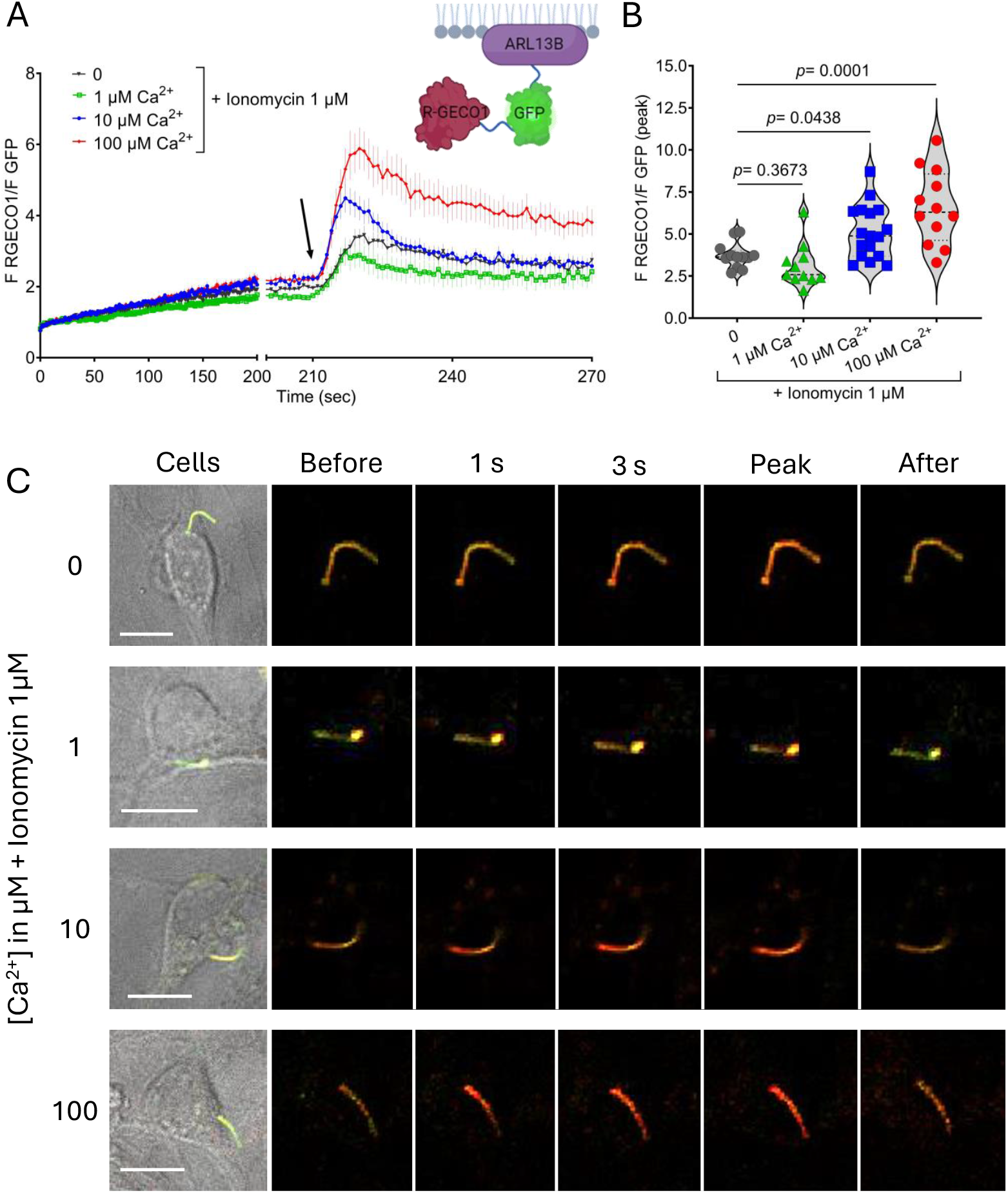
The cilia targeted calcium sensor ARL13B-GFP-RGECO1. On DIV 2-3, hippocampal cultured cells were transduced overnight with the lentivector Arl13B-GFP-RGECO1 (MOI= 1-2) and 5 days after transduction, ratiometric experiments were performed under a confocal microscope. Ciliated neurons were then filmed (1 frame/sec) before and after the application of solutions containing different calcium concentrations (0, 1, 10 or 100 μM) in the presence of ionomycin (1 μM). Fluorescence intensity was calculated by the ratio of RGEGO1 (red)/GFP (green) and normalized by the fluorescence average of the initial 30 sec of filming. (**A**) The time course of fluorescence intensity before and after solutions application. (**B**) The peak of fluorescence intensity of each solution evaluated. One-way ANOVA followed by Holm-Sidak’s multiple comparisons test was performed. p values are described in the figure (**C**) Representation of the ciliated neurons chosen for the experiment and the fluorescence course before and after each calcium solutions application. Scale bar: 10 μm.

### 3.3 Cilia-targeted Ca^2+^ and cAMP sensors coupled to the 5-HT6R in neurons

Hippocampal neuron primary cilia extend from the soma and form serotonergic synapses with axons that house GPCR effort that control the epigenetic signaling^15^. Common polymorphisms in serotonin receptor type 6 (5-HTR6) are associated with early onset of Alzheimer’s disease, development of schizophrenia and bipolar disorders, suggesting their role in cognition, learning, and mood regulation^18,21,46,47^. 5-HT6R is highly expressed brain regions like the hippocampus, cortex, and basal ganglia^18^ but these receptors almost exclusively traffic to the primary cilia of neurons. 5-HT6R is a GPCRs coupled to Gαs pathway that activates adenylyl cyclase which catalyzes production of cAMP from ATP, but it may also stimulate a non-canonical Gαq_/11_-Rho pathway in cilium-axon synapse^15^. To assess local [Ca^2+^] and [cAMP] changes in primary cilia using the same cilia-targeted moiety, we coupled 5-HT6R to RGECO1 (5-HT6R-RGECO1) within a lentivector and compared these results with the commercially available BacMan 5-HT6R-cADDi cilia cAMP sensor (**Table 2**). Like the ARL13B-GFP-RGECO1 sensor, validation experiments of the 5-HT6R-RGECO1 sensor demonstrates Ca^2+^-dependent sensitivity after permeabilized by ionomycin (**Figure 4A, B, Supplemental Video 3**). We used a pharmacological approach to validate the 5-HT6R-cADD primary cilia cAMP sensor. Here, cAMP-dependent fluorescence increased upon WAY 181187 (5-HT6R agonist) application, but this effect was ablated when neurons were incubated with the 5-HT6R antagonist, SB 271046 (**Figure 4C, D, Supplemental Figure 3**). To simultaneously assess [Ca^2+^] and [cAMP] changes within the primary cilia, we co-expressed 5-HT6R-RGECO1 with 5-HT6R-cADDis in the cultured primary hippocampal neurons (**Figure 4, Supplemental Video 4**). Stimulation of cAMP production by WAY 181187 was detected by the 5-HT6R-cADDis sensor, whereas the Ca^2+^-dependent 5-HT6R-RGECO1 fluorescence response remained unchanged. However, Ca^2+^ permeabilization by ionomycin only enhanced the 5-HT6R-RGECO1 fluorescence, whereas the signal from the cAMP sensor remained unchanged. These observations demonstrate the fidelity of using these probes in reporting unambiguous fluorescence levels proportional to [cAMP] and [Ca^2+^] measured within the same primary cilia compartment. In addition, our results indicate primary cilia [Ca^2+^] and GPCR-mediated [cAMP] dynamics are not interdependent, a finding which delineates this organelle’s localized signaling from the Ca^2+^-cAMP interplay reported globally the soma^48^. Clearly, these findings highlight the importance of using the cilia-specific sensors described when testing signal transduction pathways within this privileged organelle.

**Figure 4:**
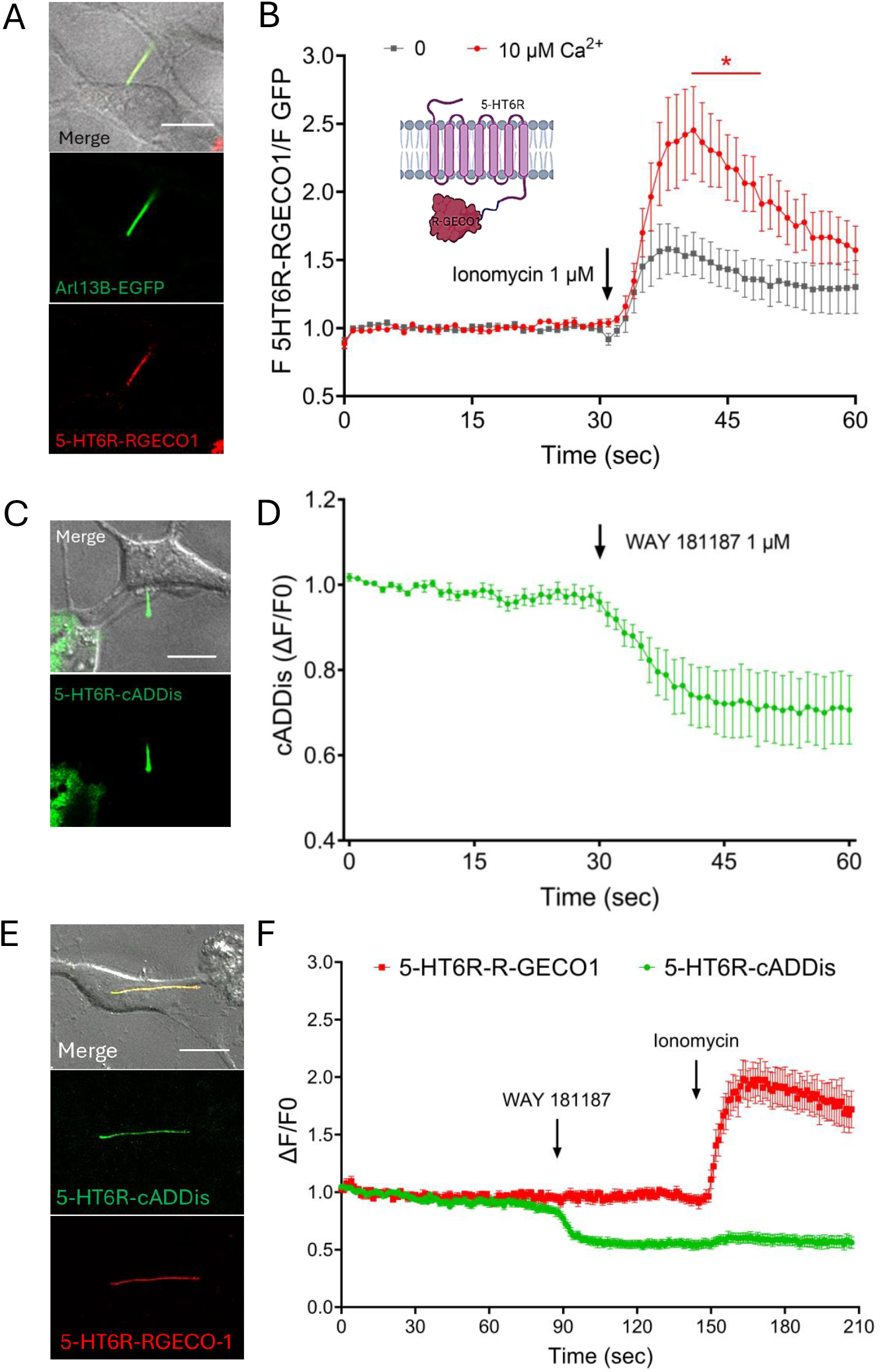
Combining primary cilia sensors for simultaneous Ca^2+^ and cAMP measurements. 5-HT6R-RGECO1 alone or in combination with the cilia cAMP sensor 5-HT6R-cADDis. Hippocampi was dissected from P0 Arl13B-EGFP^tg^ (**A, B**) or WT (**C-F**) mice and cells were seeded in glass bottom dishes. On DIV 2-3, hippocampal cultured cells were transduced overnight with the lentivector 5-HT6R-RGECO1 (MOI= 1-2) and 5 days after transduction, ratiometric experiments were performed under a confocal microscope. For the 5-HT6R-cADDis, cells were transduced one day before the experiments. Ciliated neurons were then filmed (1 frame/sec) before and after the application of solutions containing ionomycin alone (1 μM) (n= 9) or with calcium (10 μM) (n= 9) (**B**), 5-HT6R agonist WAY 181187 (1 μM) (n= 5) (**D**), or with both WAY 181187 (1 μM) followed by ionomycin (1 μM) + Ca^2+^ (10 μM) (n= 13) (F). Fluorescence intensity was calculated by the ratio of RGEGO1 (red)/GFP (green) and normalized by the F0 (fluorescence average of the initial 30 sec of filming) (**B**) or by the ΔF/F0 (**D, F**). Scale bar: 10 μm. Unpaired t test for B, * p< 0.05.

## 4. SUMMARY

We have described in detail methodologies and validation of fluorescence sensors for the detection cAMP and Ca^2+^ within hippocampal neuronal primary cilia. Previous work has reported the use of similar sensors in fibroblast, epithelial and pancreatic beta immortalized cell lines (**Table 1**)^36-38,49^. Thus to our knowledge, this is the first methods report of their use in primary hippocampal neurons. In addition we have provided details of the electrophysiology approach for directly recording PKD2L1 in the primary cilia— a Ca^2+^ channel which regulates hippocampal interneuron excitability^6^. It is expected that neuronal primary cilia also express other ion channels that conduct monovalent cations (K^+^) and anions (Cl^-^) to maintain its ionic homeostasis and the organelles resting membrane potential. Thus, the described primary cilia patch clamp method can be used to define other cilia ion channel properties. Although not presented here, the Ca^2+^ or cAMP sensors could be combined with electrophysiology to assess the impact of GPCR modulation of channel function. Recent work has expanded the tools available to include light-activated GPCRs to assess neuronal cilia regulation of behavior in vivo^50^ and nanobody-directed targeting of optogenetic tools to control cAMP levels in the primary cilium^51,52^. Take together these expanding methodologies and experimental approaches can be used to uncover underlying molecular dysfunction and signaling that cause neuronal ciliopathies; electrochemical signaling across the axo-ciliary synapse; and illuminate the basic biology of primary cilium.

## Acknowledgments

LFK was funded by the Sao Paulo Research Foundation (FAPESP, 2019/05882-8, 2019/26414-2). P.G.D. was supported by the National Institute of Diabetes and Digestive and Kidney Diseases (R01 DK123463-01, R01 DK131118-01) and the PKD Foundation (Research Grant).

## Materials availability statement

All mammalian cell expression constructs used in this study are available without restriction upon written request to the corresponding author or otherwise will be made available through Addgene (https://www.addgene.org/).

## Data Availability Statement

Primary data sets and analysis are archived through Northwestern University Institutional Repository (ARCH) and will be made available upon publication of this manuscript (DOI: https://doi.org/10.21985/n2-s825-ay98).

## FIGURE LEGENDS

**Supplemental Figure 1:**
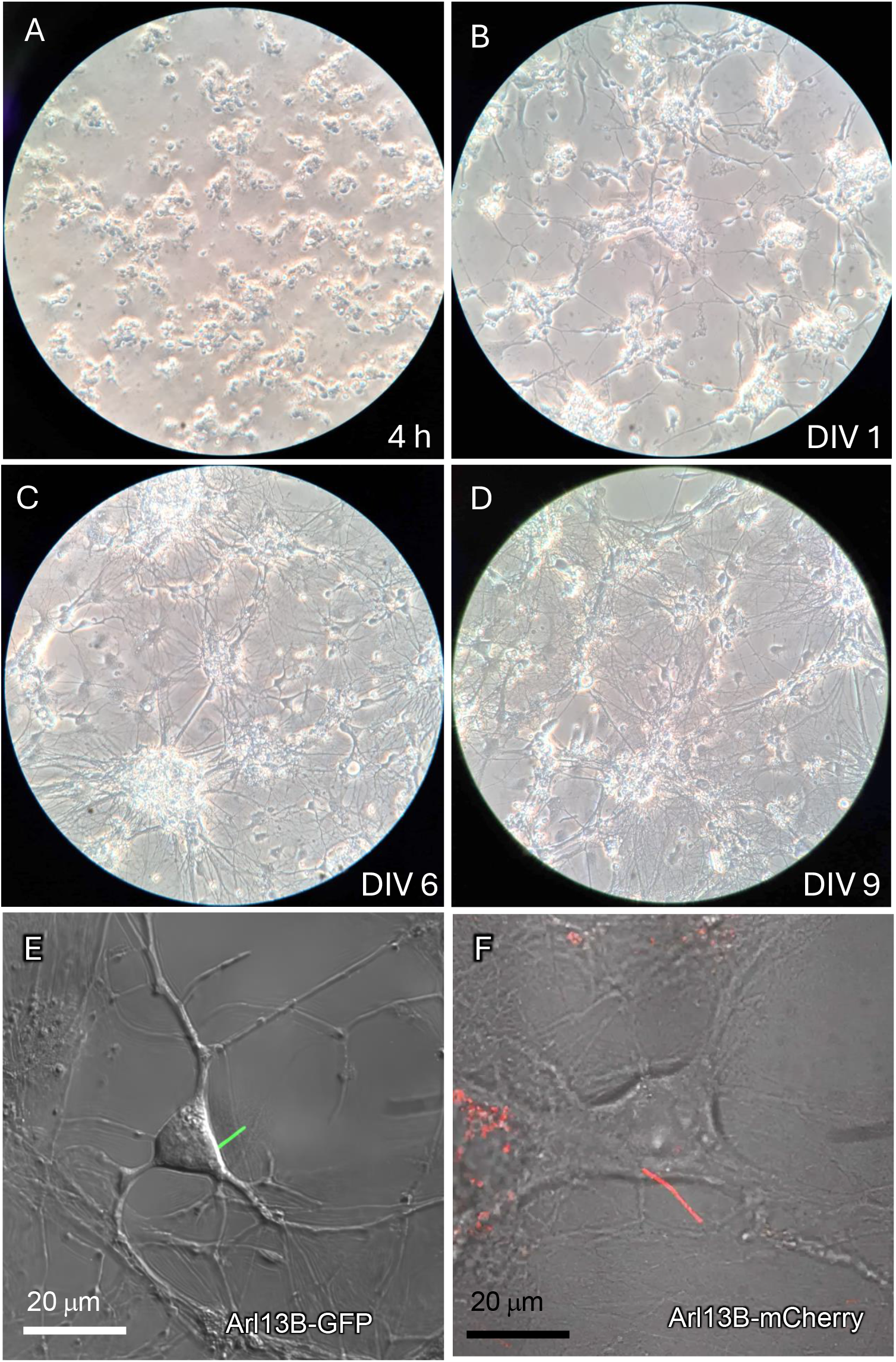
Cultures of ciliated neonatal hippocampal neurons. Hippocampi were dissected from P0 mice (n= 6-8), dissociated, cell-strained and cells were seeded in glass bottom dishes (1×10^5^ cells/dish). Images were taken under a light microscope (40x) close to the border of the glass where the density of cells are not too high and allows better visualization of the primary cilia within time. (**A**) Cells after 4 h of seeding, (**B**) DIV 1, (**C**) DIV 6, and (**D**) DIV 9. (**E, F**) Cultured hippocampal neurons at DIV 6 after infection with lentivirus for primary cilia identity reporters ARL13B-EGFP or ARL13B-mCherry.

**Supplemental Figure 2:**
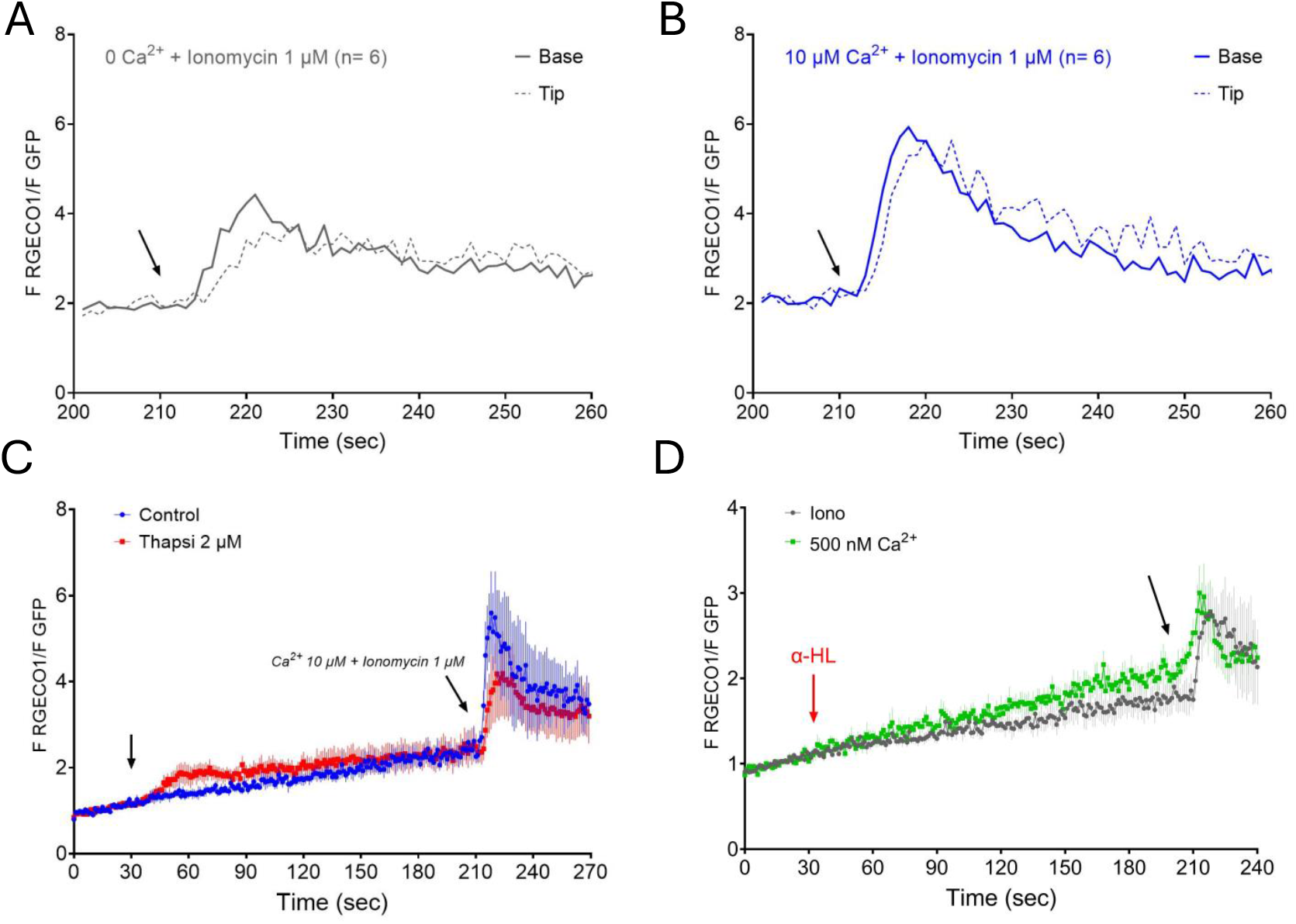
Ionomycin favors Ca^2+^ influx of from the cytoplasm to the primary cilia by releasing calcium from cytoplasmatic compartments. On DIV 2-3, hippocampal WT cultured cells were transduced overnight with the lentivector ARL13B-GFP-RGECO1 (MOI= 1-2) and 5 days after transduction, ratiometric experiments were performed under a confocal microscope. Ciliated neurons were then filmed (1 frame/sec) before and after the application of ionomycin alone (1 μM) (**A**) or in a combination with Ca^2+^ (10 μM) (**B**). ROIs were placed on both base and tip of the cilia. To verify the role of stored calcium in the influx to the primary cilia, thapsigargin (Thapsi, 2 μM, blue, n= 5) or DMSO (control, red, n= 5) treatments were performed for 3 min followed by the application of ionomycin together with Ca^2+^ 10 μM. Interestingly, we observed a small increase in calcium influx in the cilia seconds after thapsigargin application. After 3 min, the sensor response to ionomycin together with Ca^2+^ 10 μM in the cilia was smaller when compared to control group (DMSO) (**C**). When cells were permeabilized with α-hemolysin (α-HL, 10 μg/mL) for 3 min, the sensor also detected calcium changes in the primary cilia upon external application of 500 nM calcium (n= 9). Ionomycin 1 μM was used as control (n= 6) (D). Fluorescence intensity was calculated by the ratio of RGEGO1 (red)/GFP (green) and normalized by the fluorescence average of the initial 30 sec of filming.

**Supplemental Figure 3:**
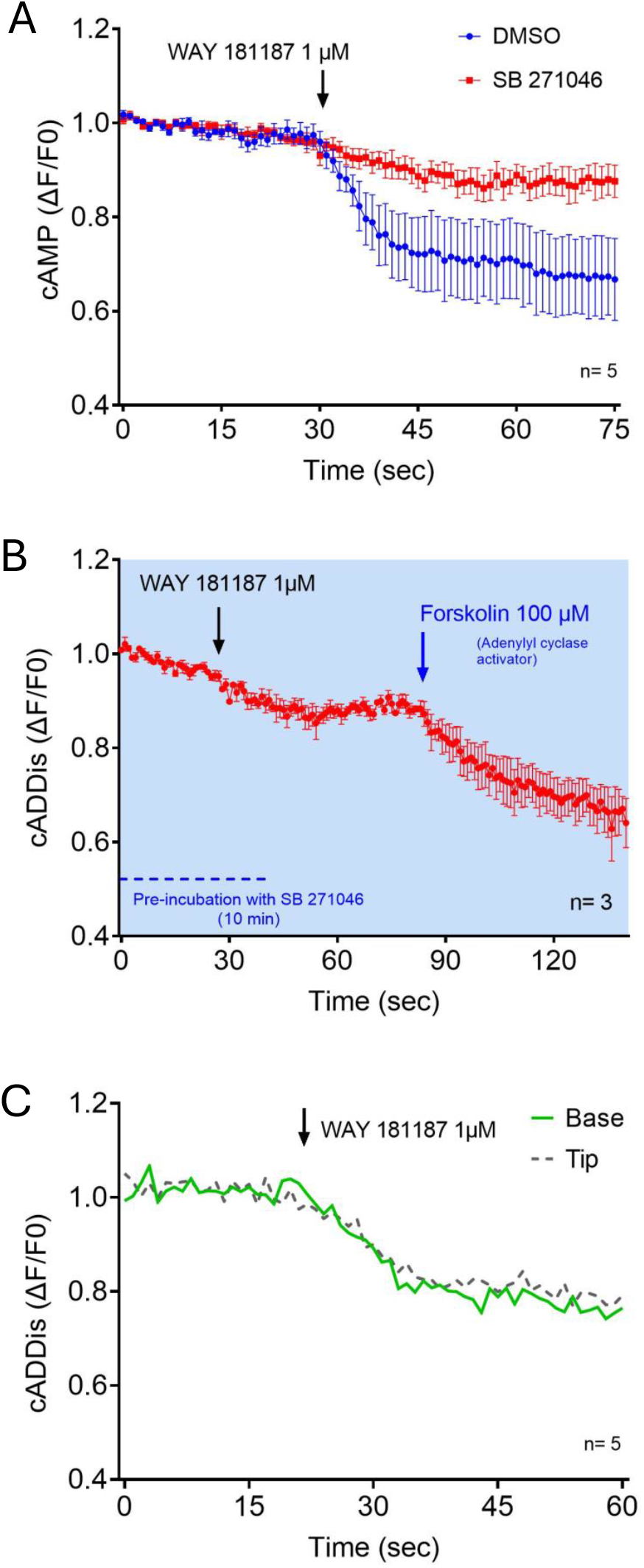
Specificity of cAMP sensor when expressed in neuronal primary cilia. On DIV 5, hippocampal WT cultured cells were transduced overnight with 5-HT6R-cADDis and 1 day after transduction, fluorescence experiments were performed under a confocal microscope. Ciliated neurons were then filmed (1 frame/sec) after the pre-incubation with SB 271046 (100 μM, n= 5) or DMSO (control, n=5) followed by the 5-HT6R agonist WAY 181187 (**A**). To confirm receptor specificity to the primary cilia 5-HT6R, we applied WAY 181187 followed by forskolin (100 μM, an activator of adenylyl cyclase) on the neurons pre-incubated with SB 271046 (100 μM) (n= 3) (**B**). To understand if cAMP production was coming from the cytoplasmic 5-HT6R rather than ciliary 5-HT6R, ROIs were placed on both base and tip of each cilium analyzed. As can be observed, cAMP production induced by WAY 181187 was homogeneous on both base and tip, indicating that cAMP is being produced within the primary cilia (**C**).

**Supplemental Video 1. Primary cilia endogenous Ca**^**2+**^ **waves**.

**Supplemental Video 2. Primary cilia ratiometric Ca**^**2+**^ **sensor (ARL13B-GFP-RGECO1) responses to ionomycin with Ca**^**2+**^ **100 μM**.

**Supplemental Video 3. Primary cilia Ca**^**2+**^ **sensor (5-HT6R-RGECO1) responses to ionomycin with Ca**^**2+**^ **10 μM**.

**Supplemental Video 4. Fluorescence responses from cAMP (5-HT6R-cADDis) and Ca**^**2+**^ **(5-HT6R-RGECO1) sensors co-expressed in hippocampal primary cilia**.

## REFERENCES

1 Pazour, G. J. & Witman, G. B. The vertebrate primary cilium is a sensory organelle. Curr Opin Cell Biol 15, 105–110, doi:10.1016/s0955-0674(02)00012-1 (2003)

2 Jurisch-Yaksi, N., Wachten, D. & Gopalakrishnan, J. The neuronal cilium - a highly diverse and dynamic organelle involved in sensory detection and neuromodulation. Trends Neurosci 47, 383–394, doi:10.1016/j.tins.2024.03.004 (2024)

3 Gopalakrishnan, J. et al. Emerging principles of primary cilia dynamics in controlling tissue organization and function. EMBO J 42, e113891, doi:10.15252/embj.2023113891 (2023)PMC10620770.

4 Plotnikova, O. V., Pugacheva, E. N. & Golemis, E. A. Primary cilia and the cell cycle. Methods Cell Biol 94, 137–160, doi:10.1016/S0091-679X(08)94007-3 (2009)PMC2852269.

5 Macarelli, V., Leventea, E. & Merkle, F. T. Regulation of the length of neuronal primary cilia and its potential effects on signalling. Trends Cell Biol 33, 979–990, doi:10.1016/j.tcb.2023.05.005 (2023)PMC7615206.

6 Vien, T. N., Ta, M. C., Kimura, L. F., Onay, T. & DeCaen, P. G. Primary cilia TRP channel regulates hippocampal excitability. Proc Natl Acad Sci U S A 120, e2219686120, doi:10.1073/pnas.2219686120 (2023)PMC10235993.

7 Ott, C. M. et al. Ultrastructural differences impact cilia shape and external exposure across cell classes in the visual cortex. Curr Biol 34, 2418–2433 e2414, doi:10.1016/j.cub.2024.04.043 (2024)PMC11217952.

8 Wu, J. Y. et al. Mapping of neuronal and glial primary cilia contactome and connectome in the human cerebral cortex. Neuron 112, 41–55 e43, doi:10.1016/j.neuron.2023.09.032 (2024)PMC10841524.

9 Klink, B. U., Gatsogiannis, C., Hofnagel, O., Wittinghofer, A. & Raunser, S. Structure of the human BBSome core complex. Elife 9, doi:10.7554/eLife.53910 (2020)PMC7018512.

10 Tian, X., Zhao, H. & Zhou, J. Organization, functions, and mechanisms of the BBSome in development, ciliopathies, and beyond. Elife 12, doi:10.7554/eLife.87623 (2023)PMC10356136.

11 Bangs, F. & Anderson, K. V. Primary Cilia and Mammalian Hedgehog Signaling. Cold Spring Harb Perspect Biol 9, doi:10.1101/cshperspect.a028175 (2017)PMC5411695.

12 Niehrs, C., Da Silva, F. & Seidl, C. Cilia as Wnt signaling organelles. Trends Cell Biol, doi:10.1016/j.tcb.2024.04.001 (2024)

13 Breunig, J. J. et al. Primary cilia regulate hippocampal neurogenesis by mediating sonic hedgehog signaling. Proc Natl Acad Sci U S A 105, 13127–13132, doi:10.1073/pnas.0804558105 (2008)PMC2529104.

14 Park, S. M., Jang, H. J. & Lee, J. H. Roles of Primary Cilia in the Developing Brain. Front Cell Neurosci 13, 218, doi:10.3389/fncel.2019.00218 (2019)PMC6527876.

15 Sheu, S. H. et al. A serotonergic axon-cilium synapse drives nuclear signaling to alter chromatin accessibility. Cell 185, 3390–3407 e3318, doi:10.1016/j.cell.2022.07.026 (2022)PMC9789380.

16 Schulz, S., Handel, M., Schreff, M., Schmidt, H. & Hollt, V. Localization of five somatostatin receptors in the rat central nervous system using subtype-specific antibodies. J Physiol Paris 94, 259–264, doi:10.1016/s0928-4257(00)00212-6 (2000)

17 Cone, R. D. Anatomy and regulation of the central melanocortin system. Nat Neurosci 8, 571–578, doi:10.1038/nn1455 (2005)

18 Dupuy, V. et al. Spatiotemporal dynamics of 5-HT(6) receptor ciliary localization during mouse brain development. Neurobiol Dis 176, 105949, doi:10.1016/j.nbd.2022.105949 (2023)

19 Barbeito, P. & Garcia-Gonzalo, F. R. HTR6 and SSTR3 targeting to primary cilia. Biochem Soc Trans 49, 79–91, doi:10.1042/BST20191005 (2021)

20 Siljee, J. E. et al. Subcellular localization of MC4R with ADCY3 at neuronal primary cilia underlies a common pathway for genetic predisposition to obesity. Nat Genet 50, 180–185, doi:10.1038/s41588-017-0020-9 (2018)PMC5805646.

21 Wachten, D. & Mick, D. U. Signal transduction in primary cilia - analyzing and manipulating GPCR and second messenger signaling. Pharmacol Ther 224, 107836, doi:10.1016/j.pharmthera.2021.107836 (2021)

22 Handel, M. et al. Selective targeting of somatostatin receptor 3 to neuronal cilia. Neuroscience 89, 909–926, doi:10.1016/s0306-4522(98)00354-6 (1999)

23 Lee, J. E. & Gleeson, J. G. Cilia in the nervous system: linking cilia function and neurodevelopmental disorders. Curr Opin Neurol 24, 98–105, doi:10.1097/WCO.0b013e3283444d05 (2011)PMC3984876.

24 Valente, E. M., Rosti, R. O., Gibbs, E. & Gleeson, J. G. Primary cilia in neurodevelopmental disorders. Nat Rev Neurol 10, 27–36, doi:10.1038/nrneurol.2013.247 (2014)PMC3989897.

25 Mill, P., Christensen, S. T. & Pedersen, L. B. Primary cilia as dynamic and diverse signalling hubs in development and disease. Nat Rev Genet 24, 421–441, doi:10.1038/s41576-023-00587-9 (2023)PMC7615029.

26 Waters, A. M. & Beales, P. L. Ciliopathies: an expanding disease spectrum. Pediatr Nephrol 26, 1039–1056, doi:10.1007/s00467-010-1731-7 (2011)3098370.

27 Guo, J. et al. Developmental disruptions underlying brain abnormalities in ciliopathies. Nat Commun 6, 7857, doi:10.1038/ncomms8857 (2015)PMC4515781.

28 Roosing, S. et al. Mutations in CEP120 cause Joubert syndrome as well as complex ciliopathy phenotypes. Journal of medical genetics 53, 608–615, doi:10.1136/jmedgenet-2016-103832 (2016)5013089.

29 Oh, E. C. & Katsanis, N. Cilia in vertebrate development and disease. Development 139, 443–448, doi:10.1242/dev.050054 (2012)3275291.

30 Sattar, S. & Gleeson, J. G. The ciliopathies in neuronal development: a clinical approach to investigation of Joubert syndrome and Joubert syndrome-related disorders. Dev Med Child Neurol 53, 793–798, doi:10.1111/j.1469-8749.2011.04021.x (2011)PMC3984879.

31 Harris, P. C. & Torres, V. E. Genetic mechanisms and signaling pathways in autosomal dominant polycystic kidney disease. The Journal of clinical investigation 124, 2315–2324, doi:10.1172/JCI72272 (2014)4089452.

32 Chebib, F. T., Sussman, C. R., Wang, X., Harris, P. C. & Torres, V. E. Vasopressin and disruption of calcium signalling in polycystic kidney disease. Nature reviews. Nephrology 11, 451–464, doi:10.1038/nrneph.2015.39 (2015)4539141.

33 Mirra, V., Werner, C. & Santamaria, F. Primary Ciliary Dyskinesia: An Update on Clinical Aspects, Genetics, Diagnosis, and Future Treatment Strategies. Frontiers in pediatrics 5, 135, doi:10.3389/fped.2017.00135 (2017)5465251.

34 Helou, J. et al. Mutation analysis of NPHP6/CEP290 in patients with Joubert syndrome and Senior-Loken syndrome. J Med Genet 44, 657–663, doi:10.1136/jmg.2007.052027 (2007)PMC2597962.

35 Ta, C. M., Vien, T. N., Ng, L. C. T. & DeCaen, P. G. Structure and function of polycystin channels in primary cilia. Cell Signal 72, 109626, doi:10.1016/j.cellsig.2020.109626 (2020)PMC7373203.

36 Su, S. et al. Genetically encoded calcium indicator illuminates calcium dynamics in primary cilia. Nat Methods 10, 1105–1107, doi:10.1038/nmeth.2647 (2013)PMC3860264.

37 Delling, M., DeCaen, P. G., Doerner, J. F., Febvay, S. & Clapham, D. E. Primary cilia are specialized calcium signalling organelles. Nature 504, 311–314, doi:10.1038/nature12833 (2013)4112737.

38 Jiang, J. Y., Falcone, J. L., Curci, S. & Hofer, A. M. Direct visualization of cAMP signaling in primary cilia reveals up-regulation of ciliary GPCR activity following Hedgehog activation. Proc Natl Acad Sci U S A 116, 12066–12071, doi:10.1073/pnas.1819730116 (2019)PMC6575585.

39 Pazour, G. J., San Agustin, J., Follit, J. A., Rosenbaum, J. L. & Witman, G. B. Polycystin-2 localizes to kidney cilia and the ciliary level is elevated in orpk mice with polycystic kidney disease. Curr Biol 12, R378–380, doi:10.1016/s0960-9822(02)00877-1 (2002)

40 Kleene, S. J. & Kleene, N. K. The native TRPP2-dependent channel of murine renal primary cilia. Am J Physiol Renal Physiol 312, F96–F108, doi:10.1152/ajprenal.00272.2016 (2017)5283891.

41 DeCaen, P. G., Delling, M., Vien, T. N. & Clapham, D. E. Direct recording and molecular identification of the calcium channel of primary cilia. Nature 504, 315–318, doi:10.1038/nature12832 (2013)PMC4073646.

42 Liu, X. et al. Polycystin-2 is an essential ion channel subunit in the primary cilium of the renal collecting duct epithelium. Elife, doi:10.1101/215814 (2017)

43 Neher, E. & Sakmann, B. Noise analysis of drug induced voltage clamp currents in denervated frog muscle fibres. J Physiol 258, 705–729, doi:10.1113/jphysiol.1976.sp011442 (1976)PMC1309001.

44 Cramer, T. M. L. & Tyagarajan, S. K. Protocol for the culturing of primary hippocampal mouse neurons for functional in vitro studies. STAR Protoc 5, 102991, doi:10.1016/j.xpro.2024.102991 (2024)PMC11017351.

45 Liu, C. & Hermann, T. E. Characterization of ionomycin as a calcium ionophore. J Biol Chem 253, 5892–5894 (1978)

46 Babic Leko, M. et al. Serotonin Receptor Gene Polymorphisms Are Associated with Cerebrospinal Fluid, Genetic, and Neuropsychological Biomarkers of Alzheimer’s Disease. Biomedicines 10, doi:10.3390/biomedicines10123118 (2022)PMC9775360.

47 Chiu, H. J. et al. Serotonin 6 receptor polymorphism in schizophrenia: frequency, age at onset and cognitive function. Neuropsychobiology 43, 113–116, doi:10.1159/000054876 (2001)

48 Halls, M. L. & Cooper, D. M. Regulation by Ca2+-signaling pathways of adenylyl cyclases. Cold Spring Harb Perspect Biol 3, a004143, doi:10.1101/cshperspect.a004143 (2011)PMC3003456.

49 Sanchez, G. M. et al. The beta-cell primary cilium is an autonomous Ca2+ compartment for paracrine GABA signaling. J Cell Biol 222, doi:10.1083/jcb.202108101 (2023)PMC9813980.

50 Winans, A. M. et al. Ciliary localization of a light-activated neuronal GPCR shapes behavior. Proc Natl Acad Sci U S A 120, e2311131120, doi:10.1073/pnas.2311131120 (2023)PMC10614621.

51 Hansen, J. N. et al. Nanobody-directed targeting of optogenetic tools to study signaling in the primary cilium. Elife 9, doi:10.7554/eLife.57907 (2020)PMC7338050.

52 Truong, M. E. et al. Vertebrate cells differentially interpret ciliary and extraciliary cAMP. Cell 184, 2911–2926 e2918, doi:10.1016/j.cell.2021.04.002 (2021)PMC8450001.

